# Modelling metastasis in zebrafish unveils regulatory interactions of cancer-associated fibroblasts with circulating tumour cells

**DOI:** 10.1101/2022.12.07.519426

**Authors:** Pablo Hurtado, Inés Martínez-Pena, Sabrina Yepes-Rodríguez, Miguel Bascoy-Otero, Carmen Abuín, Cristóbal Fernández-Santiago, Laura Sánchez, Rafael López-López, Roberto Piñeiro

## Abstract

The dynamic intercommunication between tumour cells and cells from the microenvironment, such as cancer-associated fibroblast (CAFs), is a key factor driving breast cancer (BC) metastasis. Clusters of circulating tumour cells (CTCs), known to bare a higher efficiency at establishing metastases, are found in the blood of BC patients, often accompanied by CAFs in heterotypic CTC-clusters. Previously we have shown the utility of CTC-clusters models and the zebrafish embryo as a model of metastasis to understand the biology of breast cancer CTC-clusters. In this work, we use the zebrafish embryo to study the interactions between CTCs in homotypic clusters and CTC-CAFs in heterotypic CTC-clusters to identify potential pro-metastatic traits derived from CTC-CAF communication. We found that upon dissemination CAFs seem to exert a pro-survival and pro-proliferative effect on the CTCs, but only when CTCs and CAFs remain joined as cell clusters. Our data indicate that the clustering of CTC and CAF allows the establishment of physical interactions that when maintained over time favour the selection of CTCs with a higher capacity to survive and proliferate upon dissemination. Importantly, this effect seems to be dependent on the survival of disseminated CAFs and was not observed in the presence of normal fibroblasts. Moreover, we show that CAFs can exert regulatory effects on the CTCs without being involved in promoting tumour cell invasion, and these effects are differential based on the BC cell molecular phenotype, and the crosstalk between tumour cells and CAFs, i.e. paracrine or physical interaction. Lastly, we show that the physical communication between BC cells and CAFs leads to the production of soluble factors involved in BC cell survival and proliferation. These findings suggest the existence of a CAF-regulatory effect on CTC survival and proliferation sustained by cell-to-cell contacts and highlight the need to understand the molecular mechanisms that mediate the interaction between the CTCs and CAFs in clusters enhancing the metastatic capacity of CTCs.

## 1 Introduction

Circulating tumour cells (CTCs) are thought to be the tumour cell populations that give rise to distant metastases. CTCs can travel as either individual cells or as small oligoclonal groups of cells known as CTC-clusters (1). These CTC populations have clinical relevance, and their presence in the blood of cancer patients is associated with poor disease outcomes in several cancer types, such as breast, prostate, and colorectal cancer (1–7). Despite the low occurrence of CTC-clusters in the bloodstream of cancer patients (1), they have been shown to have a higher metastatic potential than individual CTCs due to factors such as the existence of heterogeneous CTC populations, cell-cell junctions enhancing resistance to anoikis, decreased expression of genes associated with apoptosis and immune response, and increased expression of genes associated with proliferation and stemness (8–12). Therefore, CTC-clusters seem to play a prominent role in metastasis, and broadening our knowledge of their biology may be key to designing promising therapeutic strategies against metastasis.

Cancer progression is supported by the dynamic crosstalk among cancer cells and other cellular components that infiltrate the tumour such as immune cells, vascular cells, inflammatory cells, and fibroblasts. This communication influences disease initiation, progression, organ-specific metastasis, and patient prognosis (13, 14). Thus, it has become clear that tumour progression does not exclusively rely on cancer cell-autonomous functions and that tumour stroma reactivity is a key contributing factor. Indeed, in the context of metastatic spread, the analysis of the blood of patients and cancer animal models has revealed the presence of stromal cells – endothelial cells, platelets, fibroblasts, leukocytes, and neutrophils – along with CTCs, predominantly forming the known as heterotypic CTC-clusters (15). Among these stromal cells, tumour-activated fibroblasts (aka. cancer-associated fibroblasts, CAFs) are known to contribute to cancer initiation, progression, and metastasis (16, 17). Thus, CAFs are involved in processes such as tumourigenesis, angiogenesis (18), immunosuppression (19), drug resistance (20), maintenance of cancer stemness, and metabolic reprogramming (21–23).

CAFs have been identified as circulating cells in the blood of patients with breast, prostate, lung, and colorectal cancer, and they can cluster with CTCs (24, 25). Particularly in breast cancer (BC), the dissemination of heterotypic CTC-clusters seems to be an early event, as they are found in all clinical stages of BC (26). In addition, CAFs are also found not only at the primary site but also at distant organs, suggesting that they may support the homing and growth of disseminated CTCs (27–29). Moreover, the interaction of CTCs with CAFs seems to exert an enhancement on the survival and metastatic competence of CTCs (24, 26, 30). It has been proposed that the pluripotency marker CD44 may be mediating heterotypic interactions between populations of BC CTCs and CAFs. In addition, homophylic CD44 interactions promote CTC-clustering and the development of polyclonal metastases in BC models, and the presence of CD44+ CTC-clusters correlates with a poor prognosis for patients with BC (8). This evidence indicates that the mechanisms involved in CTC-CAF clustering may be important in the regulation of cell-to-cell communication and enhanced CTC survival and metastases seeding of CTC-clusters. However, little is yet known about how they work and to what extent are dependent on physical CTC-CAF contact or paracrine interactions through the exchange of soluble factors.

We previously reported that CTC-clusters have intrinsic properties derived from the clustering of tumour cells that represent a metastatic advantage over single CTCs. Making use of the zebrafish and mouse xenografts as *in vivo* models of BC metastasis, we showed that disseminated CTC-clusters bare a higher survival and proliferation capacity than single CTCs, due to increased resistance to fluid shear stress (FSS), and supported by the upregulation of genes involved in cell cycle and stemness (31). Building up on this knowledge, in the present work we set out to investigate the role of the interactions between CTCs and CAFs in heterotypic CTC-clusters to identify potential pro-metastatic traits of these circulating entities. Surprisingly, the injection into zebrafish of heterotypic CTC-clusters, derived from the co-culture of BC cell lines (MDA-MB-231 and MCF7) and a CAF cell line (vCAF), or homotypic CTC-clusters did not show significant differences on the invasive capacity of CTCs, suggesting that not all CAF populations are capable of driving invasión. Importantly, our data indicate that not all disseminated CTCs and CAFs remain as clusters, but when they do, the presence of CAFs seems to exert a pro-survival and pro-proliferative effect on the CTCs. This effect seems to be CAF-specific, as normal fibroblasts do not affect disseminated CTC numbers, and dependent on the survival and physical interaction of CAFs over time. Lastly, molecular analyses show that the communication between BC cells and CAFs is dependent on physical interactions, deriving in the production of soluble factors involved in tumour cell survival and proliferation such as RANTES and CXCL10.

## 2 Materials and methods

### 2.1 Cell culture and generation of single and CTC-cluster *in vitro* models

*In vitro* homotypic CTC-clusters were generated using the human triple-negative MDA-MB-231 and the luminal A MCF7 BC cell lines, and heterotypic CTC-clusters were generated in combination with the vCAF fibroblast, from the vulvar carcinoma CAFs61 fibroblasts, or the healthy fibroblasts BJ cells, from the human foreskin, in a 2:1 ratio (tumour cell: fibroblast). Tumour cell lines stably express the enhanced green fluorescent protein (eGFP) and the luciferase gene, while vCAF are labelled with the red fluorescent protein (mCherry). For the visualization of BJ cells, fluorescent lipidic stains were used. MDA-MB-231 and MCF7 were purchased from GeneCopoeia, Inc (USA), vCAF were provided by Dr Erik Sahai, and BJ cells were obtained from the American Type Culture Collection (ATCC). MDA-MB-231, MCF7 and BJ cells were cultured in DMEM High Glucose supplemented with 10% fetal bovine serum (FBS) and 1% Penicillin/Streptomycin (P/S), at 37 °C in a humidified atmosphere containing 5% CO_2_. In addition, the vCAF culture medium was supplemented with Insulin + Transferrin + Selenium (GIBCO).

According to the experimental needs, two different protocols were used for the generation of single and CTC-cluster models: i) The required number of cells was cultured overnight in low-adherence conditions to allow cell aggregation. The day after, cell suspensions were physically dissociated by gentle pipetting to obtain a single-cell suspension and a small cell-group suspension, corresponding to single CTC and CTC-cluster models, respectively; ii) Adherent cell monolayers at an 80% confluence were dissociated with different concentrations trypsin-EDTA (LONZA), 10X (0,25% v/v) and 1X (0,05% v/v), to obtain either an individual cell suspension or a clustered-cell suspension, respectively.

### 2.2 Cell viability and proliferation

Tumour cell proliferation was measured through the detection of luciferase activity. Homotypic and heterotypic cultures were seeded in a 96-well opaque plate (2.5 x 10^3^ cells/well) in 200 μL of cell culture medium. Cells were incubated for 4, 24, 48 and 72 hours, D-luciferin (Perkin Elmer) was added to a final concentration of 150 μL/mL and luciferase signal was measured using the Envision® 2105 Multimode Plate Reader (Perkin Elmer). Alternatively, AlamarBlue^®^ proliferation assay was performed. Cells were incubated for 4, 24, 48 and 72 hours and 10% AlamarBlue^®^ was added, measuring the fluorescent signal after a 3 hour incubation using FLUOstar OPTIMA (BMG Labtech) plate reader.

### 2.3 Transwell® migration

Migration capacity of homotypic and heterotypic cultures was measured using 6.5 mm Transwell® with 8.0 μm Pore Polycarbonate Membrane Insert (Corning, Madrid, Spain). The transwells were coated with 100 μL of growth factor reduced Matrigel (10 ug/μL) and incubated for 30 minutes enabling polymerization. MDA-MB-231 or MCF7 were co-cultured with vCAF 24 hours before the assay to obtain heterotypic clusters and were maintained in serodeprivation. The day after, 5 × 10^4^ cells/ Transwell® were seeded in a 10% FBS gradient (37°C, 5% CO_2_) for 8 hours (MDA-MB-231) or 24 hours (MCF7). After incubation, the insert was washed with phosphate saline buffer (PBS) (BIOWEST), and the top area of the insert was scraped to remove the non-migrated cells. Migrated cells were fixed with 3% paraformaldehyde (PFA) + 1% glutaraldehyde (Sigma-Aldrich) for 15 minutes and GFP positive cells were counted using Leica DMi8 (Leica Microsystems) and the free software ImageJ Fiji (NIH Image).

### 2.4 Wound healing

MDA-MB-231 or MCF7 (2 × 10^5^ cells) were seeded in a 24 wells plate. Once an 80% confluence monolayer was obtained, vCAF^mCherry^ cells were added and cultures incubated at 37 °C for 24 hours, allowing a complete monolayer formation. A wound was created in the monolayer, the culture medium was retired, the wells washed with PBS to eliminate dead cells, and a fresh medium was added. Images were obtained at 0, 20 and 28 hours (MDA-MB-231) or 0, 24 and 48 (MCF7) using a Leica Dmi8 microscope (Leica Microsystems), and wound size was observed over time. Wound width was determined by measuring micrometres between two parts of monolayer in the same location at every time point with IamgeJ Fiji (NIH Image).

### 2.5 Soft-agar colony formation assay

Homotypic or heterotypic clusters (1 x 10^4^ tumor cells/well) were resuspended in 0.3 % (w/v) molecular grade agarose (Fisher BioReagents, Madrid, Spain) in DMEM (10% FBS, 1% P/S) inside a 6 wells plate. Cells were incubated at 37°C, 5% CO_2_ for 3-4 weeks to allow colonies to spawn, At the end of the experiment, colonies were fixed with 3% PFA + 1% glutaraldehyde (Sigma-Aldrich) for 15 minutes. The number and area of colonies were measured using a Leica DMi8 microscope (Leica Microsystems) and the free software ImageJ Fiji (NIH Image). Because of the small size of MDA-MB-231 colonies, a minimum size of 75 μm was established to quantify colonies from MDA-MB-231 homotypic and heterotypic clusters.

### 2.6 Gene expression analysis

The analysis of gene expression was performed by RT-qPCR. RNA was extracted using the High Pure Isolation Kit (Roche Life Science, Barcelona, Spain), following manufacturer instructions, and later quantified using a Nanodrop 2000 spectrophotometer (Thermo Fisher Scientific, Waltham, MA, USA). The reverse transcription process (RT-PCR) was performed using the SuperScript^™^ III Reverse Transcriptase (Thermo Fisher Scientific), according to the specifications of the manufacturer: 25 °C for 5 minutes, 50 °C for 2 hours and 70 °C for 15 minutes. cDNA was amplified through the use of TaqMan^®^ probes (Supplementary Table S1, Thermo Fisher Scientific) and TaqMan^®^ Universal PCR Master Mix (Life Technologies, Carlsbad, CA, USA) in a LightCycler^®^ system (Roche Life Science, Barcelona, Spain). PCR amplification was performed following these steps: denaturation at 95 °C for 10 minutes, 45 cycles of 95 °C for 10 seconds, 60 °C for 30 seconds, and 72 °C for 10 seconds. Gene expression levels were normalized by ΔCt, using average expression of glyceraldehyde-3-phosphate dehydrogenase gene (GAPDH) and β-2-Microglobulin (B2M). Each sample was processed in duplicate.

### 2.7 Spheroid formation and invasion assay

Heterotypic suspensions of MDA-MB-231 and vCAF (4 x 10^4^) were cultured as spheroids by the hanging drop method. Cells were dissociated from a monolayer, counted and resuspended in MammoCult medium (StemCell Technologies Cat# 05620) at a concentration of 5 x 10^4^ cells/ml. Fourty aliquiots (20μL each) of the suspension (containing 1000 cells each) were deposited on the underside of a 10 cm petri dish lid. The lid was then inverted over the dish and filled with 10 ml of PBS. The dish was maintained at 37°C in a humidified incubator with 5% CO2 for 7 – 14 days, until the spheroids were visible at naked eye. Spheroids were then harvested in an Eppendorf tube and allowed to settle at the bottom. After, 40μL of spheroids were carefully aspirated and mixed with 100μL of matrigel and 100μL of collagen and deposited in a 24-well plate previously coated with matrigel. The plates were incubated at 37°C for one hour and then 1 mL of complete DMEM medium was added. Pictures were taken at 0, 24 and 48 hours using a Leica Microsystems DMi8 fluorescence microscope.

### 2.8 Transforming growth factor β1 (TGF-β) treatment

vCAF (6 x 10^4^) or BJ (1 x 10^5^) cells were seeded in a 12 well plate and incubated at 37 °C with DMEM (10% FBS, 1% P/S) for 48 hours. Cultures were serum-starved in DMEM (1% P/S) for 24 hours, and the human TGF-β (5 ng/mL) was added (Peprotech, Spain) for 48 hours. TGF-β was reconstituted with citric acid, which was used as a control in this assay. Wells were washed with PBS and lysis buffer from High Pure RNA Isolation Kit, to obtain the RNA (section Gene expression analysis).

### 2.9 Fluid Shear Stress (FSS) Assay

Cell suspensions (5 × 10^5^ cells/mL in a total volume of 4 mL) were loaded into a 5 mL syringe (NORM-JECT) attached to a 30-gauge needle. The syringes were loaded into a syringe pump (Harvard Apparatus, MA, USA), which generates shear forces in order to mimic the mechanical stress that tumor cells undergo into the bloodstream. It was considered one passage (P) when the entire content of the syringe passed through the 30-gauge needle. Each cell suspension was passed through the needle at a flow rate of 4.45 mL/min (1840 dyn/cm^2^) for a total of 10 passages, with a period of 2 min resting between passages. Aliquots of 100 μL were taken from the cell suspensions before the first passage (P0), as well as after even passages (2nd, 4th, 6th, 8th, and 10th) to assess cell viability, right after finishing the assay (0 h) and after culturing the cells for 24 h. Cell viability was determined by incubating the cell with a luciferin solution (150 μg/mL; Perkin Elmer, MA, USA) and quantification of the bioluminescent signal emitted in an EnVision plate reader (Perkin Elmer).

### 2.10 Immunofluorescence

Cells (1 x 10^4^) were seeded over lenses in a 24 wells plate and incubated with DMEM (10% FBS, 1% P/S) at 37 °C for 24 hours. After that, the medium was retired, wells were washed with PBS and cells were fixed with PFA 4% for 15 minutes. Wells were washed 3 times with PBS and cells were blocked with 10% goat serum (Sigma-Aldrich) for 30 minutes. Insideperm (Inside Stain Kit, Miltenyi Biotec) was used as permeabilizer and antibody preparation, and the primary antibodies against α-SMA (Ab7817; Abcam) and FAP (Ab53066; Abcam) were incubated for 1 hour at room temperature or overnight at 4 °C. After incubation with the primary antibodies, cells were washed 3 times with PBS for 5 minutes and incubated with AlexaFluor^®^ 647 conjugated secondary antibodies (Ab150079, Ab150115; Abcam) for 1 hour at room temperature. Cell nuclei were stained with DAPI (4’,6-diamidino-2-phenylindole). After incubation with secondary antibody, cells were washed 3 times with PBS and fixed to a slide using 5 μL of Mowiol (4-88 Reagent, Calbiochem). Fluorescent images were acquired with a Leica Dmi8 microscope (Leica Microsystems).

### 2.11 Cytokine detection assay

Cytokine levels produced by homotypic and heterotypic cultures were analyzed using the Proteome Profiler^™^ Human Cytokine Array (ARY005B; R&D SYSTEMS) kit. MDA-MB-231 or vCAF cultures were seeded in different plates, and MDA-MB-231 + vCAF cocultures were seeded in the same plate, allowing physical interactions, or separated by 0.4 μM pore Transwell (Corning), to study paracrine interactions. The number of cells seeded was adjusted to allow all cultures to reach 80% confluence at the same time when the FBS was retired and cells were incubated with DMEM (P/S) for 48 hours. Supernatants were retired and centrifuged at 264 xg for 5 minutes to eliminate suspension particles. Samples were processed following manufacturer instructions and incubated over membranes. These membranes were revealed at 40 seconds of exposure using a Chemidoc MP system (Bio-Rad) and cytokine expression levels were determined by densitometry using the free software ImageJ Fiji (NIH Image).

### 2.12 Zebrafish (*Danio rerio*) embryo xenograft

Zebrafish embryos were generated through the mating of adult zebrafish. Embryos were maintained at 28 °C and incubated with PTU 0,003% weight/volume (N-Phenylthiourea, Sigma) in E3 medium, to inhibit superficial pigmentation. Two days old larvae were dechorionated and anesthetised with tricaine 0,003 % weight/volume in PTU (Sigma), and placed in a 10 cm Petri plate covered with a 1,5 % agarose layer. Right before the microinjection, cellular suspensions were obtained from cultures and 1 x 10^6^ million tumour cells were resuspended in 6 μL of 2% PVP (Polivinilpirrolidin, Sigma-Aldrich). Microinjection needles were crafted from borosilicate capillaries (1mm outer diameter x 0,58 mm inner diameter, Harvard Apparatus) and the needle tip was segmented at 0,7 cm from the beginning of the narrowing. Approximately, 250 cells were injected into the perivitelline space of embryos using a microinjector IM300 (Narishige) with an output pressure of 10 psi and a time injection of 0,3 ms. After injection, the fluorescence signal was observed in the injection location and embryos were incubated in a 96 wells plate with PTU at 34 °C for 3 days (120 hours post fertilization). Images of every embryo were obtained using a fluorescence microscope DMi8 (Leica Microsystems) at 0, 24, and 72 hours to determine cellular dissemination, survival, and proliferation. Moreover, colocalization between disseminated tumour cells (eGFP) and disseminated vCAFs (mCherry) was analyzed through fluorescence overlapping. BJ cells were labelled with 5 μL per million cells of Vibrant^™^ DiD Cell-Labelling Solution (TermoFisher Scientific), a lipidic fluorescent staining that allows cell visualization at 647 wavelength, and incubated for 5 minutes at 37 °C and 15 minutes at 4 °C to avoid colorant internalization. Data are representative of more than 30 embryos in 3 or more assays. The minimum embryo survival rate in the analyzed assays is 70%.

### 2.13 Mouse (*Mus musculus*) lung colonization assay

Female Scid-Beige mice from the Barcelona Biomedical Research Park (PRBB) were kept in pathogen-free conditions at the animal facility of the University of Santiago de Compostela. The experimental procedure was approved by the Animal Experimentation Ethical Committee of the University of Santiago de Compostela (15010/2019/002) and all animal experiments complied with the EU Directive 2010/63/EU for animal experiments. Homotypic MDA-MB-231 (5 x 10^5^ cells) cell clusters and heterotypic MDA-MB-231 + vCAF (5 x 10^5^ + 5 x 10^5^) clusters were resuspended in 100 μL of PBS and injected into the lateral tail vein of 10 weeks old SCID-beige mouse. Mice were immobilized with a restraining system (trap) and cells were injected using 27 gauge needles attached to a 1 mL syringe. One hour after injection, tumor cell presence in the lungs was determined using the *in vivo* image system IVIS (Perkin Elmer) after the intraperitoneal injection of luciferin (150 mg of luciferin/kg; XenoLight^™^ D-Luciferin K+ Salt Bioluminescent, Perkin Elmer). Metastasis development was monitored every 3-4 days until 18 days when mice were sacrificed by CO_2_ inhalation and their lungs were extracted. Lungs were fixed with 10% w/v formaldehyde (VWR) and embedded in paraffin. Lung cuts were stained with hematoxylin, eosin and antibodies for human keratins AE1/AE3 (760-2135; Roche diagnostics). Metastasis quantification was reached through a review of 17 digitalized slides. We used 3DHISTECH’s SlideViewer version 2.5 (3DHISTECH Ltd. Budapest, Hungary). During this process, each metastatic focus was associated with a marker from the Marker Counter tool, which allowed us to obtain the total marker count and, thus, the total number of metastases per slide.

### 2.14 Data analysis

Microscopic images from in vitro and in vivo assays were analyzed with Leica Application Suite X software (Leica Microsystems) and free ImageJ Fiji software (NIH Image). Fluorescence images from zebrafish xenografts were analyzed using the free software Quantifish (Zebrafish Image Analyser). Quantification, statistic analysis and graphic representation were performed using the GraphPad Prism 8 software (GraphPad Software). Data from functional in vitro assays, and mouse and zebrafish xenografts, when different experimental groups are compared, were statistically analyzed using the Mann-Whitney test. Zebrafish experiments, when different time points were compared in one experimental group, were analyzed using the Wilcoxon test (non-parametric test for paired samples). Regarding the correlation study in zebrafish, a linear regression analysis was performed.

## 3 Results

### 3.1 Establishment and functional characterization of heterotypic CTC-clusters of BC tumour cell lines and CAFs

To investigate the effects of CAFs on BC cells with different intrinsic phenotypes the cell lines MCF7 and MDA-MB-231, stably expressing eGFP-luciferase, were co-cultured with the CAF cell line vCAF, expressing mCherry, and a series of *in vitro* assays were carried out to evaluate the functional characteristics involved in the different phases of the metastatic process. In co-culture conditions, vCAF cells slowed down the proliferation and wound closure ability of MDA-MB-231 cells, but increased the proliferation and wound closure ability of MCF7 cells (Figure 1A, B). In addition, monocultures of BC cells and co-cultures with vCAF fibroblasts were established to generate homotypic (^hom^CTC-C) and heterotypic (^het^CTC-C) clusters. Under co-culture conditions, both BC cell lines were able to efficiently form heterotypic clusters with the fibroblasts (Supplementary Figure S1A). When the colonogenesis capacity of clusters was tested in soft agar experiments, ^het^CTC-C produced a similar number of colonies as ^hom^CTC-C (Figure 1C), although showed a significantly smaller colony diameter when formed by MDA-MB-231 cells, which was not observed for clusters of MCF7 cells (Figure 1C). Interestingly, ^het^CTC-C from both BC lines displayed an enhanced migratory capacity, suggesting that vCAF fibroblast might increase the invasive potential of BC cell lines in the transwell assay (Figure 1D). On the other hand, co-cultures established under paracrine conditions - separated by a permeant membrane allowing the exchange of soluble factors - did not induce significant changes in the mRNA levels of CDH1, VIM, and MKI67 in the BC cells, genes associated with epithelial-to-mesenchymal transition (EMT) and proliferation (Supplementary Figure S1B-D), neither in the expression of genes associated with CAF biology, such as FN1, FSP1, ACTA2, FAP, CDH2 and VIM (Supplementary Figure S1E-J).

**Figure 1.**
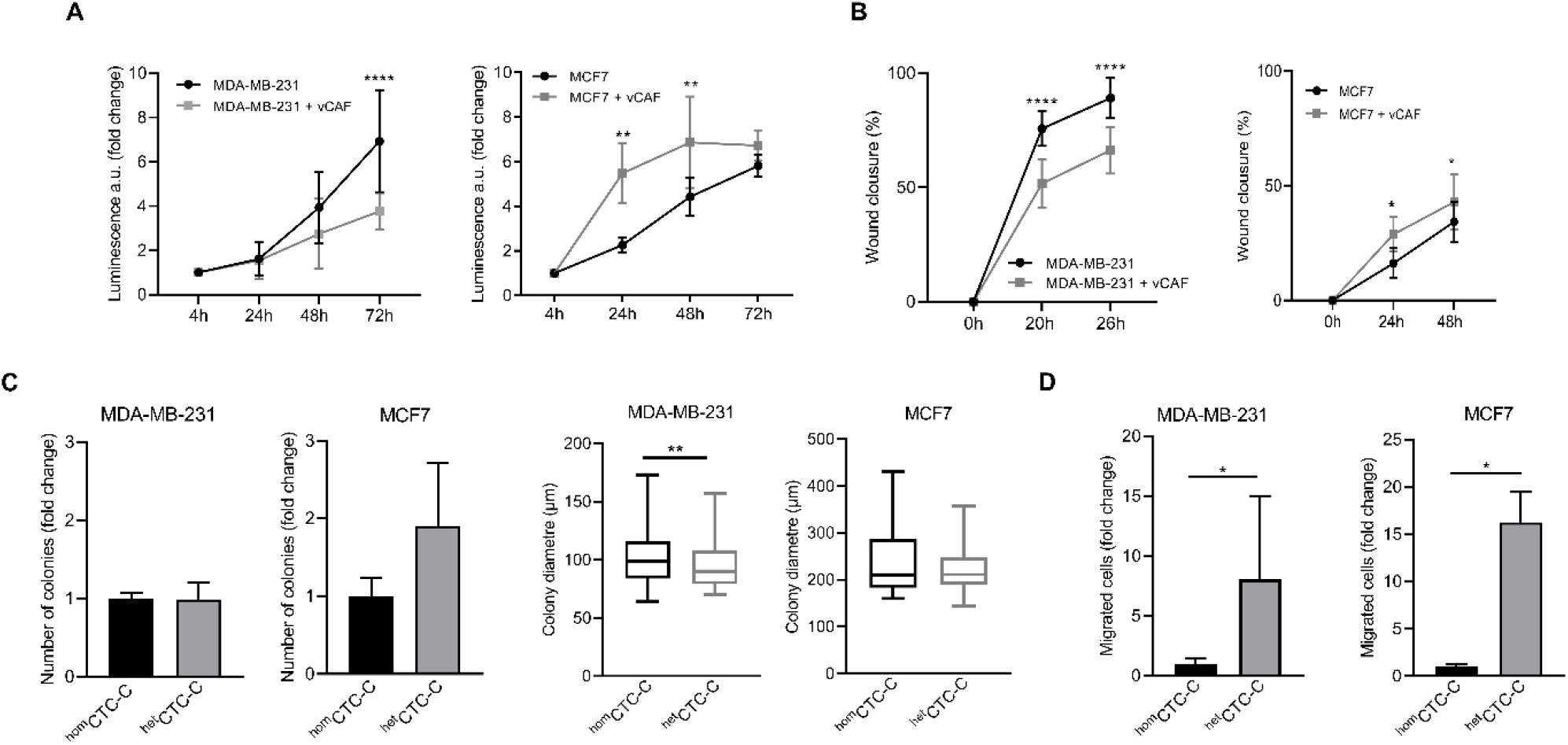
Evaluation of the effect of vCAF fibroblasts on BC cell lines by *in vitro* functional assays. **(A)** Proliferation of MDA-MB-231 (n= 3) and MCF7 (n=4) cells in presence of vCAF along 72 hours. Data are expressed as the fold change of proliferation relative to time 4 hours; **(B)** Wound healing capacity of MDA-MB-231 (n= 5) and MCF7 (n= 2) cells in presence or absence of vCAF; **(C)** Number of colonies generated by ^hom^CTC-C and ^het^CTC-C from MDA-MB-231 (n= 3) and MCF7(n=3) cells in soft agar colony formation assays represented as the fold change of ^het^CTC-C over ^hom^CTC-C, and average colony diameter; **(D)** Migration of ^hom^CTC-C and ^het^CTC-C derived from MDA-MB-231 (n= 3) and MCF7 (n= 2) cell lines. Data are expressed as the fold change of migrated cells relative to the migration observed in the ^hom^CTC-C population. \**p* < 0.05, \*\**p* < 0.01, \*\*\*\**p* < 0.0001.

### 3.2 Homotypic and heterotypic CTC-clusters have similar invasive behaviour in the zebrafish

The metastatic potential of ^hom^CTC-C and ^het^CTC-C was studied in the zebrafish. CTC-clusters were xenografted at the convergence of the Duct of Cuvier, near the pericardial space, and their dissemination is evaluated in the caudal region of the fish, based on the quantification of fluorescence (eGFP) intensity (Supplementary Figure S2A). Despite the known limited metastatic capacity of the luminal MCF7 cell line *in vivo*, we set out to determine whether the clustering with vCAF fibroblasts could favour their survival and dissemination through circulation. The dissemination of clusters towards the caudal region of the fish was quite low for ^hom^CTC-C (30 ± 11%) and ^het^CTC-C (38 ± 23%) xenografts, with both populations showing a dramatic decrease in the percentage of fish showing CTC dissemination throughout the experiment (3 ± 6% in ^hom^CTC-C xenografts and 9 ± 12% in ^het^CTC-C) (Figure 2A). These data suggest that the clustering of MCF7 cells and vCAF fibroblast does not increase the limited invasive and dissemination capacity of the BC cells. However, the quantification of the fluorescence intensity of MCF7 cells at the xenograft site showed that, despite the high mortality observed for the cells throughout the experiments, the number of surviving cells was significantly higher in those fish xenotransplanted with ^het^CTC-C, indicating a possible regulatory effect of vCAF in tumour cell survival as early as 24 hours after xenotransplantation (Figure 2B, C).

**Figure 2.**
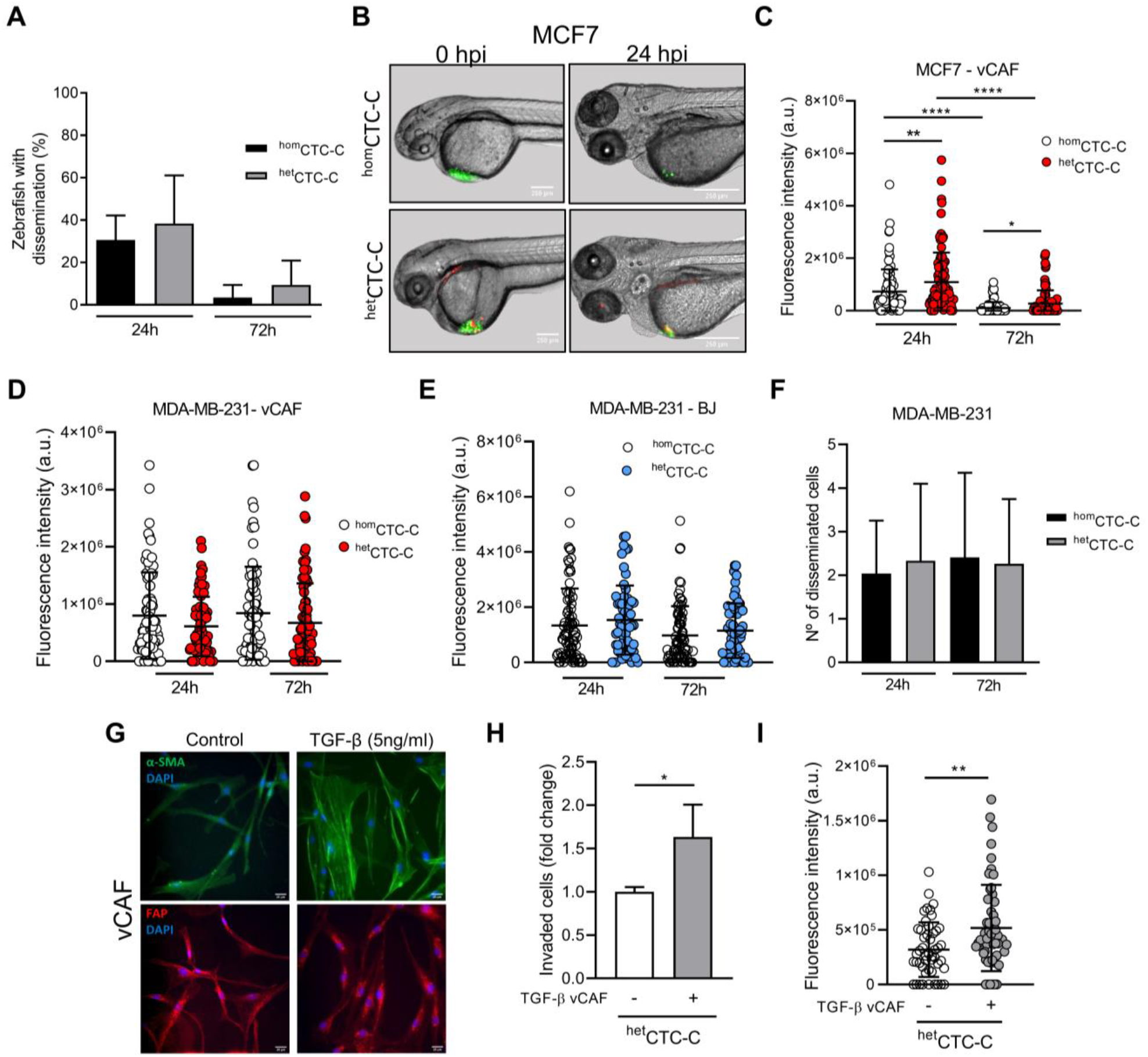
Assessing the metastatic ability of BC ^hom^CTC-C and ^het^CTC-C in the zebrafish. **(A)** Percentage of fish xenografted with MCF7 ^hom^CTC-C and ^het^CTC-C showing tumour cells dissemination in the tail 24 and 72 hpi; (**B)** Representative images of zebrafish showing ^hom^CTC-C and ^het^CTC-C from MCF7 (green) and vCAF (red) xenografts in the Duct of Cuvier right after injection (0 hpi) and 24 hpi (scale bar 250μm); **(C)** Dot plot of the fluorescence intensity (eGFP) at the injection site of fish xenografted with ^hom^CTC-C and ^het^CTC-C representing the survival of MCF7 cells. Each dot represents an individual fish (n = 3 independent experiments); **(D)** Dot plot of the fluorescence intensity at the tail of fish xenografted with MDA-MB-231 ^hom^CTC-C and vCAF ^het^CTC-C and **(E)** BJ ^het^CTC-C. Each dot represents an individual fish (n =3 independent experiments); **(F)** Number of disseminated cells found at the tail of fish xenografted with MDA-MB-231 ^hom^CTC-C and vCAF ^het^CTC-C; **(G)** Representative images of the protein expression of α-SMA (green) and FAP (red) in control and TGF-β treated vCAF. Cells were stimulated with 5 ng/mL TGF-β for 48 hours. Nuclei are labelled with DAPI (blue) (scale bar 25 μm); **(H)** Effect on the migration of MDA-MB-231 cells from ^het^CTC-C formed by untreated or TGF-β treated vCAF (n= 2); **(I)** Fluorescence intensity detected in the tails of fish xenotransplanted with ^het^CTC-C formed by untreated or TGF-β treated vCAF 24 hpi. Each dot represents an individual fish (n =1). \**p* < 0.05, \*\**p* < 0.01, \*\*\*\**p* < 0.0001.

We next studied the behavior of the highly metastatic triple negative cells MDA-MB-231 in association with vCAF. The dissemination of MDA-MB-231 CTCs showed no significant differences between ^hom^CTC-C and ^het^CTC-C, even though we observed a tendency toward higher dissemination in the fish xenotransplanted with ^hom^CTC-C, based on the fluorescence values (8 x 10^5^ ± 7.56 x 10^5^ vs. 6.12 x 10^5^ ± 5.17 x 10^5^; *p* > 0.05) (Figure 2D; Supplementary Figure S2B). Moreover, when we performed the same experiments xenografting ^het^CTC-C formed by primary fibroblast from healthy tissue (BJ) as a control, a similar result was obtained showing that the level of dissemination was the same for ^hom^CTC-C and ^het^CTC-C (Figure 2E). Altogether, these results show that vCAF fibroblast do not promote the dissemination of the highly metastatic MDA-MB-231 cells in the zebrafish.

To better characterise the invasive phenotype of the clusters, ^hom^CTC-C and ^het^CTC-C were xenotransplanted into the perivitelline space of zebrafish embryos, from where cells must actively invade and intravasate to gain access to the circulatory system (Supplementary Figure S2C). Quantification of the number of invasion fronts showed that these are significantly higher in embryos xenotransplanted with ^hom^CTC-C compared to ^het^CTC-C, both at 24 hours (2.2 ± 1 vs. 1.6 ± 1.1, p<0.05) and 72 hours (2.8 ± 1.5 vs. 2 ± 1, p<0.05) (Supplementary Figure S2D). However, no differences were observed in the number of tumour cells disseminated in the fish tails (Figure 2F). These results support the idea that the vCAF fibroblasts do not potentiate the invasive capacity of MDA-MB-231 cells and even can exert a negative effect on them. This idea was reinforced by an *in vitro* invasion assay with spheroids generated by MDA-MB-231 and vCAF showing that BC cells led the invasion fronts, followed by the fibroblasts vCAF (Supplementary Figure S2E). Furthermore, the comparative analysis of gene expression profiles of molecular markers associated with CAF activation (FN1, FSP1, ACTA2, FAP, PDGFRα, PDGFRβ, and TNC) between vCAF and BJ fibroblast showed no significant differences (Supplementary Figure S3A-G).

### 3.3 TGF-β stimulated vCAF enhance heterotypic CTC-cluster dissemination

The similar molecular profile observed for the fibroblast vCAF and BJ, together with the lack of a pro-invasive effect on the CAFs in ^het^CTC-C and the failure of MDA-MB-231 to modify the expression levels of genes associated with fibroblast activation, lead us to hypothesize that vCAF fibroblasts do not possess or are not able to acquire a phenotype promoting BC cell invasion and dissemination in the presence of BC cells. To test this hypothesis, vCAF cells were stimulated with TGF-β, a cytokine known to promote CAF activation and cancer progression (32). TGF-β treatment resulted in an increased expression of FN1 (4.2 ± 2.5 vs. 1 ± 0.3; *p* < 0.05) and FAP (6.1 ± 2.7 vs. 1 ± 0.3; *p* < 0.05), while strong trends toward increased expression of ACTA2 (1.7 ± 0.8 vs. 1 ± 0.06), TNC (3.1 ± 1.7 vs. 1 ± 0.5) and PDGFRβ (1.78 ± 0.28 vs. 1 ± 0.3) were also observed. No change was observed in the expression of FSP1 and PDGFRα (Supplementary Figure S4A-G). Moreover, gene expression changes on ACTA2 and FAP also translated into the proteins codified by these genes, α-SMA and FAP (Figure 2G). This was particularly evident for α-SMA, being incorporated into stress fibers upon TGF-β treatment. Furthermore, the invasive ability of ^het^CTC-C established with pre-stimulated vCAF was enhanced *in vitro* and in the zebrafish when compared to ^het^CTC-C of unstimulated vCAF (Figure 2H, I). This set of experiments shows that vCAF can acquire the ability to enhance tumour cell invasion in response to stimuli from the tumour environment, and may indicate the existence of inefficient crosstalk between these fibroblasts and the MDA-MB-231 cells.

### 3.4 Heterotypic CTC-clusters do not have a metastatic advantage in mice over homotypic clusters

Our zebrafish data indicates that the fibroblasts vCAF do not favour the dissemination of heterotypic CTC-clusters of MDA-MB-231 cells. To contrast these results in a mammalian and more complex model, we injected equivalent numbers of cells either as ^hom^CTC-C or ^het^CTC-C into the lateral tail vein of immunodeficient mice and measured their capacity to colonise the lungs of mice and form metastases by imaging of the bioluminescence signal of BC cells (Figure 3A). Upon cluster injection, we observed that the bioluminescence in the lungs of mice from both groups drastically decreased during the first week compared to the moment of injection (Figure 3B), which indicates CTC death due to shear stress and/or homing. The relative change in bioluminescence between day 3 after injection and day 0 was lower in mice xenotransplanted with ^hom^CTC-C than with ^het^CTC-C, indicating a better survival of the CTCs from ^hom^CTC-C (0.29 ± 0.15 vs. 0.15 ± 0.08; p> 0.05; Figure 3C), although these differences were not significant. These results suggest that ^het^CTC-C might show reduced resistance to the shear forces in the circulation compared to ^hom^CTC-C. However, when both cluster populations were subjected to high magnitude stress forces in an *in vitro* fluidic assay, ^het^CTC-C showed a similar resistance to fluid shear stress (FSS) to that of ^hom^CTC-C (Figure 3D). Only a small trend toward increased resistance of ^het^CTC-C was observed. This data goes in agreement with the zebrafish experiments, in which no significant differences were observed in the dissemination and survival of both types of clusters (Figure 2D). Furthermore, the incidence of lung metastases at the end of the experiment was significantly higher in mice injected with ^hom^CTC-C (Figure 3E), which was also confirmed by the histological examination of the lungs (^hom^CTC-C 321.3 ± 101.4 vs. ^het^CTC-C 129.4 ± 87.36; *p* < 0.05) (Figure 3F, G). Based on this evidence, we conclude that the differences observed between CTC-clusters types might be due to an impaired homing of CTCs from heterotypic clusters, perhaps due to massive cell death of vCAF fibroblasts, which may not survive in the mouse lungs. This would cause the difference in CTC survival observed at day 3 and being responsible for the delayed appearance of metastasis in the mice lungs. Indeed, the histopathological examination of the metastatic lesions of mice injected with ^het^CTC-C could not confirm the presence of vCAF.

**Figure 3.**
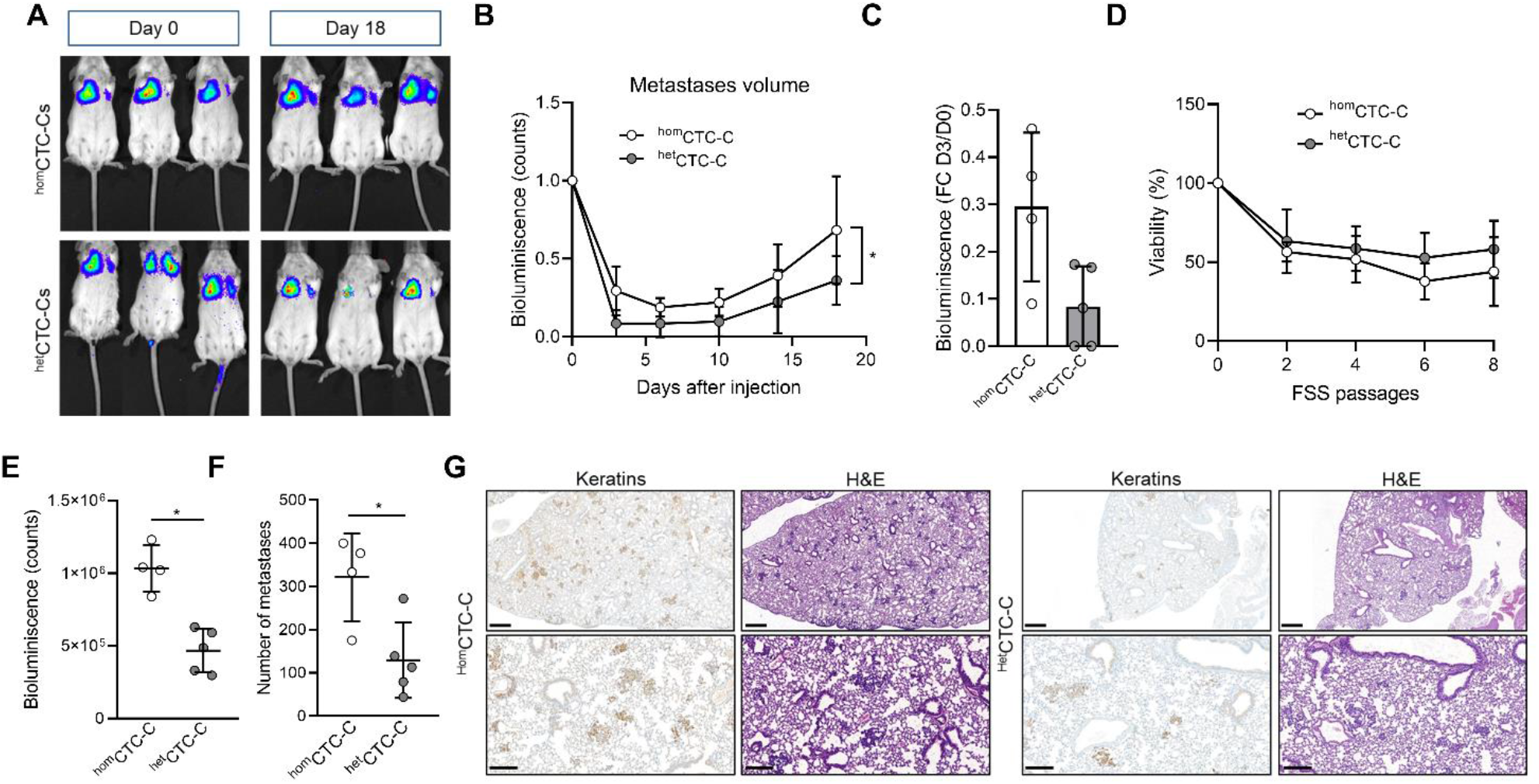
Assessing metastatic colonisation ability of ^hom^CTC-C and ^het^CTC-C in the mouse. **(A)** Representative images of the development of lung metastatic lesions over time in mice injected into the lateral tail vein with ^hom^CTC-C and ^het^CTC-C assessed by measuring the bioluminescence signal (counts); (**B)** Bioluminescence signal throughout time emitted by the tumour cells that has colonised the pulmonary tissue (at least n = 4 mice per group); **ac** Percentage of decrease of the bioluminescence signal in the lung of mice 3 days after tail vein injection of ^hom^CTC-C or ^het^CTC-C; **(D)** Cell viability of ^hom^CTC-C and ^het^CTC-C subjected to different passages of high magnitude fluid shear stress (FSS); **(E)** Quantification of the bioluminescence signal in the lungs at day 18; **(F)** Quantification of the number of metastatic foci in the lungs; **(G)** Representative images of lungs with metastases, visualized with Hematoxin-eosin (H&E), and sections of the lung stained with an antibody for keratins. Note the brown staining in tumour clusters and nodules. Images are shown at a 3x magnification (scale bar 500 μm; upper images) and a 10x magnification (scale bar 200 μm; lower images). \**p* < 0.05.

### 3.5 Disseminated CAFs promote tumour cell survival and proliferation

In contrast to the mouse as a model system, the zebrafish embryo allows for real-time visualization in of the dissemination and homing of CTCs and fibroblasts within the first days. Detailed analysis of the ^het^CTC-C xenografted embryos revealed that not all fish showed disseminated vCAF in the caudal region, although BC cell dissemination was always observed and the percentage of fish with disseminated vCAF cells decreased over time (Figure 4A). These results might indicate that the dissemination of MDA-MB-231 cells and vCAF do not always occurs as heterotypic clusters, but also that not all CAF survive upon dissemination. Based on this observation, we analysed the population of fish according to the presence or absence of disseminated vCAF. This analysis showed that the fluorescence associated with MDA-MB-231 cells, and therefore the volume of cells, was higher in the tails of fish presenting disseminated CAFs compared to those without them (Figure 4B). However, we did not find a correlation between the mCherry and eGFP fluorescence signals (R2 = 0.15 at 24h; R= 0.36 at 72h) (Supplementary Figure S5A), suggesting that the volume of disseminated BC cells is not dependent on the volume of disseminated CAFs, and vice versa. In addition, when the same experiment was performed with control fibroblasts, we observed a higher fluorescence associated with BC cells in the tail of fish showing disseminated BJ than in those without them (24h: 2. 12 x 10^6^ ± 1.3 x 10^6^ vs. 8.83 x 10^5^ ± 7.31 x 10^5^; 72h: 2.16 x 10^6^ ± 7.65 x 10^5^ vs. 9.28 x 10^5^ ± 8.93 x 10^5^; p< 0.0001) (Figure 4C), despite fish injected with ^het^CTC-C showed lower dissemination and survival of BJ fibroblasts along time (Supplementary Figure S5B). Therefore, altogether, these results seem to indicate that the co-dissemination of tumour cells and fibroblasts is not due to a vCAF-driven invasion of the BC cells.

**Figure 4.**
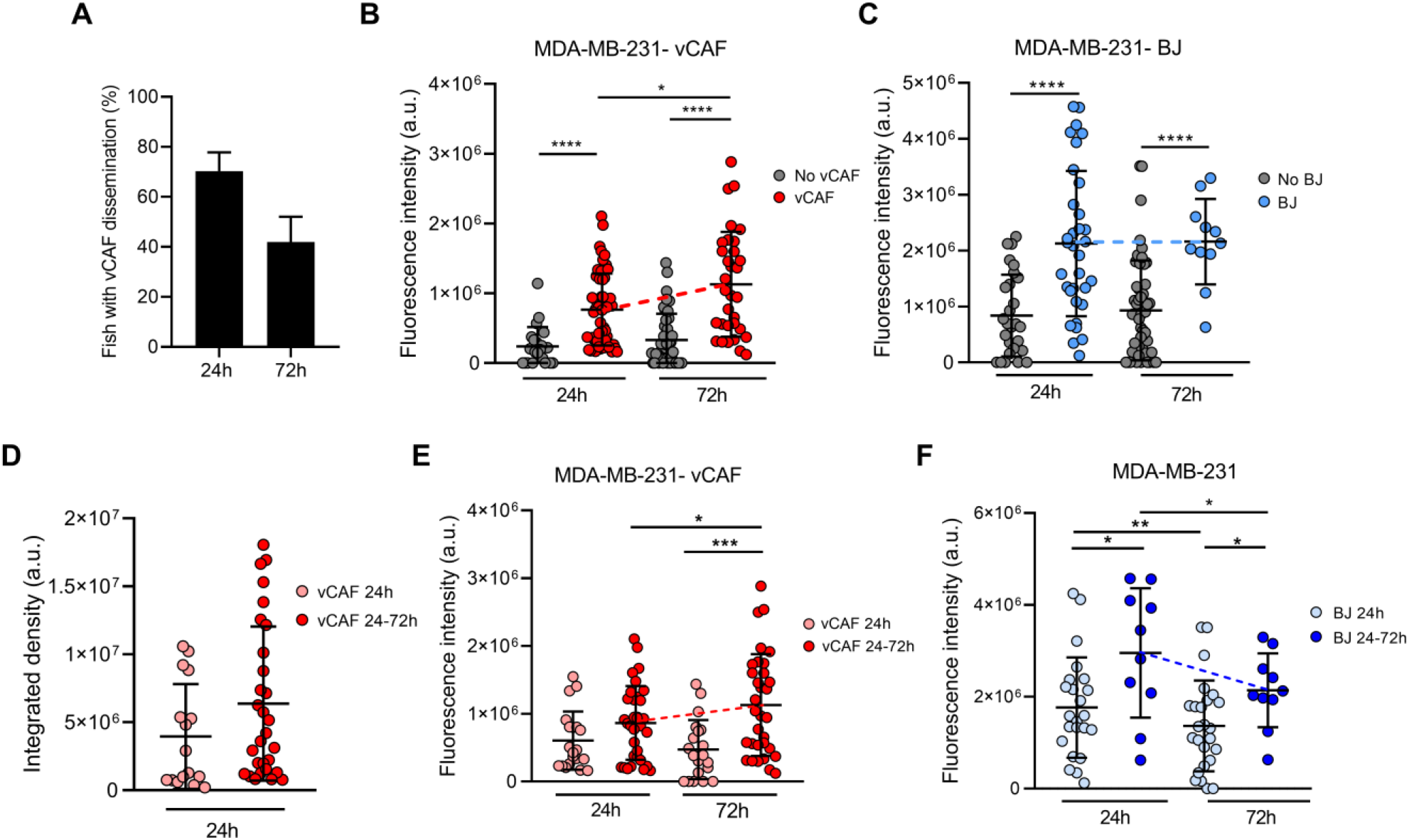
Evaluation of the survival of CTCs and CAFs in the zebrafish embryo xenografted with ^het^CTC-C. **(A)** Percentage of fish xenografted with MDA-MB-231 and vCAF ^het^CTC-C showing disseminated fibroblasts in the tail at 24 and 72 hpi; Dot plots of the fluorescence intensity of MDA-MB-231 disseminated in the fish tails according to the presence or absence of vCAF **(B)** and BJ **(C)** fibroblasts; **(D)** Dot plot of the integrated fluorescence density of vCAF disseminated in the fish tails; Dot plots of the fluorescence intensity of MDA-MB-231 disseminated in the fish tails according to the survival of vCAF **(E)** and BJ **(F)** fibroblasts throughout the experiments. Each dot represents an individual fish (n =3 independent experiments). \**p* < 0.05, \*\**p* < 0.01, \*\*\**p* < 0.001, \*\*\*\**p* < 0.0001.

Very interestingly, the experiments with vCAF ^het^CTC-C revealed a significant increase in the number of disseminated MDA-MB-231 (from 24 to 72 hpi) when the CAF were present, determined by an increase in fluorescence signal (7.6 x 10^5^ at 24h vs. 1.1 x 10^6^ at 72h; p< 0.05; Figure 4B). On the contrary, this effect was not found in embryos showing no CAF dissemination, indicating that vCAF may exert a pro-survival and pro-proliferative effect on BC cells. Importantly, and in contrast to the results observed for CAF clusters, the experiments with BJ ^het^CTC-C did not show differences in the number of BC cells between 24 and 72 hpi when BJ cells are disseminated (Figure 4C). Therefore, these results indicate that the dissemination of CTC-CAF clusters facilitates not only the survival but also the proliferation, and therefore the homing of clusters of MDA-MB-231 cells.

### 3.6 Survival of disseminated CAFs favours CTC proliferation

In order to better understand the possible regulation of vCAF on the survival and proliferation of MDA-MB-231 cells, we set out to analyse whether this effect was dependent on the sustained presence of fibroblasts over time, or it was established in the first hours after dissemination of the clusters throughout the fish circulation. Embryos with disseminated vCAF were divided and analysed in two groups according to whether they showed fibroblasts in the tail both at 24 and 72 hours (vCAF 24-72h), or only at 24 hours (vCAF 24h) after xenotransplantation. The analysis of the fluorescence signal associated with mCherry showed a trend towards a higher volume of disseminated vCAF fibroblasts at 24 hpi in those embryos of the vCAF 24-72h group compared to those vCAF 24h embryos (6.38 x 10^6^ ± 5.66 x 10^6^ vs. 3.95 x 10^6^ ± 3.86 x 10^6^; p= 0.07; Figure 4D). Follow-up over time of these fibroblasts (vCAF 24-72h) indicates that the volume of disseminated cells at 72h is not significantly different from that observed at 24h (6.27 x 10^6^ ± 6.43 x 10^6^ vs. 4.8 x 10^6^ ± 5.18 x 10^6^; p= 0.17; Supplementary Figure S5C), although a decrease associated with cell death is observed. This result would indicate that some vCAF can survive once disseminated in the fish. On the other hand, the analysis of the fluorescence signal associated with MDA-MB-231 cells revealed that the survival of vCAF throughout the experiment (vCAF 24-72h) was associated with a higher volume of MDA-MB-231 in the fish tails, compared to the CAF 24h group, and the volume of tumour cells experienced a significant increase from 24 to 72 hpi (24h = 8.4 x 10^5^ ± 5.4 x 10^5^ vs. 72h = 1.1 x 10^6^ ± 7.5 x 10^5^; p< 0.05; Figure 4E). Importantly, this result was not observed in the control experiments with the BJ fibroblasts, where despite a higher initial tumour cell volume in the BJ 24-72h group, the number of BC cells decreased significantly between the 24 and 72 hpi (2.9 x 10^6^ ± 1.4 x 10^6^ vs. 1.3 x 10^6^ ± 1.3 x 10^6^; p< 0.05; Figure 4F). Indeed, we observed that in the vCAF 24-72h group there was a higher percentage of fish in which the fluorescence signal from MDA-MB-231 cells increased from 24 to 72 hpi compared to the CAF 24h group (69.4 ± vs. 33.3 ± 16.4; p< 0.01), in contrast with the large percentage of fish in the BJ population in which the number of BC cells decreased over time (Supplementary Figure S5D, E). Taken together, these results suggest that the survival of CAFs in the zebrafish upon dissemination of heterotypic CTC-CAF clusters favours the homing of BCtumour cells, in contrast to the survival of BJ fibroblasts.

### 3.7 Physical interaction between CAFs and BC cells determines the survival and proliferation of CTCs

If the survival of fibroblasts vCAF favours the metastatic capacity of MDA-MB-231 cells, it would be expected that the integrity of heterotypic CTC-CAF clusters is preserved after dissemination and therefore fibroblasts appear in close interaction with tumour cells. The interaction between the two cell types in the fish tails was studied by the overlap between the fluorescence signals, indicative of cell colocalization and physical interaction. We found that the percentage of colocalization at 24h in the vCAF 24-72h group of fish was significantly higher than in the vCAF 24h group (8.68 ± 7.47% vs. 3.71 ± 5.29%; p< 0.01), and these percentages were maintained at 72 hpi (7.36 ± 9.12%; p> 0.05) (Figure 5A). However, this analysis represents an underestimation of the physical interaction of both cell lines, as it is only based on the determination of fluorescence overlap (Figure 5A). These results suggest that CTC-CAF direct interactions - probably due to the initial cell clustering - are a mechanism regulating the survival and proliferation of MDA-MB-231, and possibly of vCAF. This idea is strongly supported by the significant decrease observed over time in the number of disseminated BC cells when they were xenotransplanted with vCAF as heterotypic cell suspensions (^het^CTC-S) instead of clusters (Figure 5B). MDA-MB-231 cells from ^het^CTC-S had an impaired survival upon dissemination in the zebrafish, which was not previously observed for ^het^CTC-C (Figure 2D). Also, note that the survival of ^het^CTC-S was even lower than observed for ^hom^CTC-S, suggesting that vCAF exert a regulatory effect on the dissemination and survival of MDA-MB-231 cells (Figure 5B). Interestingly, the analysis of ^het^CTC-S xenografts reflected that the presence of disseminated CAFs was not accompanied by the proliferation of BC cells (Figure 5C), even when CAF survived throughout the experiment (vCAF 24-72h; Figure 5D). These results were contrary to the results obtained with ^het^CTC-C (Figure 4B), and despite the similar percentages of fish showing disseminated CAFs in the caudal region (Figure 4A and Supplementary Figure S5F). Altogether, these experiments demonstrate that the aggregation of MDA-MB-231 and vCAF in heterotypic CTC-clusters can enhance the survival and proliferation of BC CTCs by allowing the establishment of physical interactions required for the regulatory action of CAF over the metastatic capacity of CTCs.

**Figure 5.**
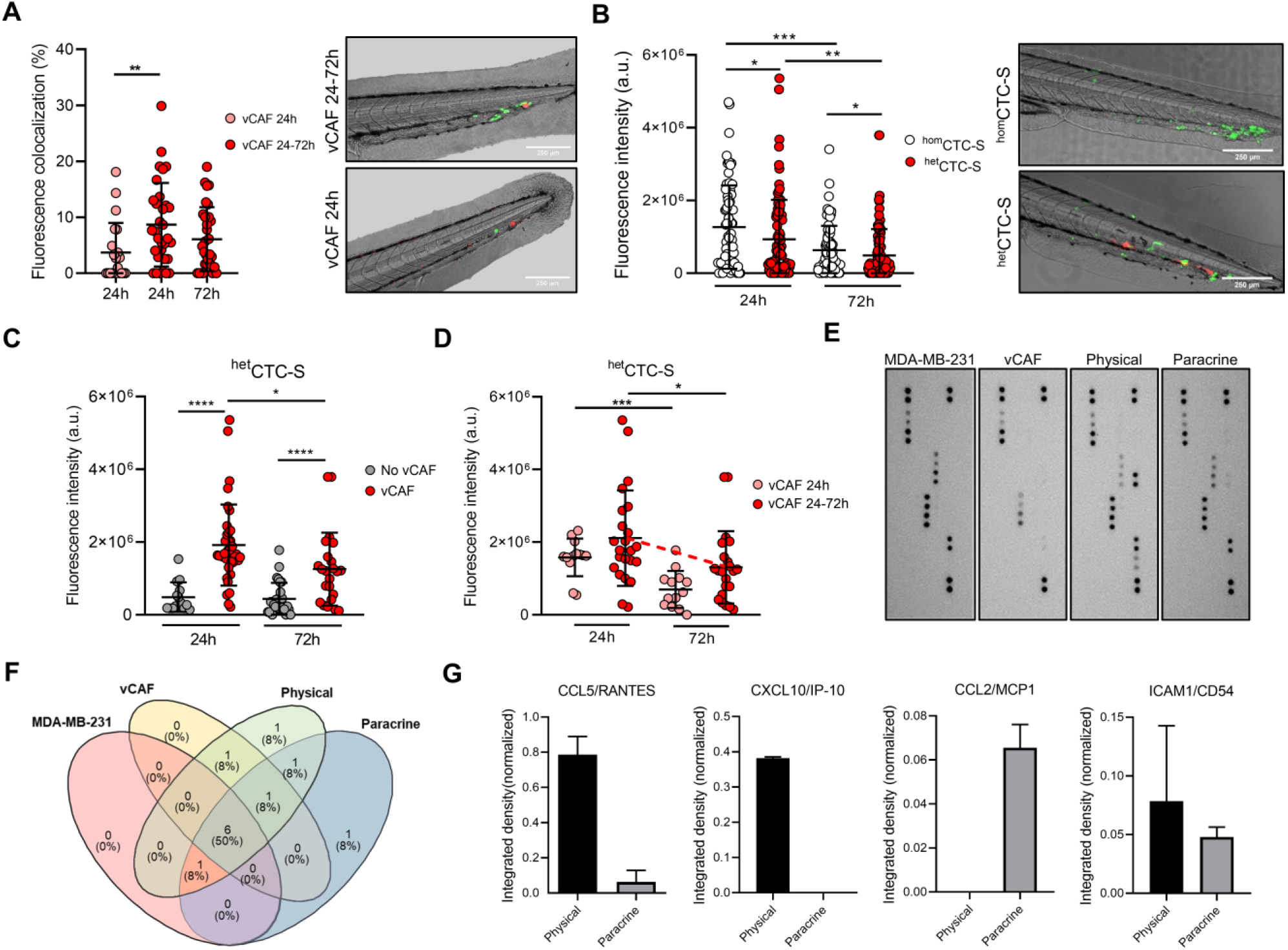
Analysis of colocalization of of CTCs and CAFs in the zebrafish embryo and cytokine profile. **(A)** Dot plot showing the percentage of fluorescence overlap between the signal emitted by the disseminated MDA-MB-231 and vCAF cells, and representative images of the tails (scale bar 250μm); **(B)** Dot plot of the fluorescence intensity in the tail of fish xenografted with MDA-MB-231 ^hom^CTC-S and vCAF ^het^CTC-S, and representative images of the tails (scale bar 250μm). Each dot represents an individual fish; **(C)** Dot plots of the fluorescence intensity of MDA-MB-231 disseminated in the fish tails xenografted with MDA-MB-231 ^hom^CTC-S and vCAF ^het^CTC-S according to the presence or absence of vCAF; **(D)** Dot plots of the fluorescence intensity of MDA-MB-231 ^het^ disseminated CTC-S in the fish tails according to the survival of vCAF; **(E)** Images of the membranes of an antibody array evaluating the production of cytokines and chemokines in the culture medium of monocultures and co-cultures of MDA-MB-231 and vCAF cells under physical and paracrine interaction. Red squares mark the spots of cytokines exclusively detected under a specific culture condition; **(F)** Venn diagram showing the cytokine/chemokine overlap among culturing conditions; **(G)** Level of cytokines/chemokines exclusively detected under physical and paracrine conditions. \**p* < 0.05, \*\**p* < 0.01, \*\*\**p* < 0.001.

To determine the molecular signals derived from the physical interactions of MDA-MB-231 and vCAF a cytokine array was done on the supernatants of cell cultures under three conditions: i) monoculture of each cell type, ii) co-culture of cells separated by a permeable membrane (paracrine interaction), and iii) co-culture allowing the direct interaction between cell types (physical interaction) (Figure 5E). Under the experimental conditions, out of the 36 cytokines evaluated by the assay (Supplementary Table S2), 12 cytokines were detected, 1 exclusively under physical interactions, 1 exclusively under paracrine interactions, and 6 cytokines were detected in all conditions (Figure 5F). Interestingly, four of the cytokines detected in co-cultures were not detected (CCL2/MCP1, CXCL10/IP-10 and ICAM1/CD54) or barely detected (CCL5/RANTES) in the monocultures (Figure 5G). CCL5/RANTES and CXCL10/IP-10 were detected when MDA-MB-231 and vCAF were cultured in physical contact, and CCL2/MCP1 only under paracrine interactions. Notably, the cytokines CCL5/RANTES, CXCL10/IP-10 and ICAM/CD54 increase tumour cell motility and invasion, with CXCL10/IP-10 and ICAM/CD54 also enhancing a metastatic phenotype in breast cancer tumour cells.(33–35) These results, together with the previously shown gene expression analyses (Supplementary Figure S1D), suggest the existence of a communication phenomenon between the MDA-MB-231 and vCAF cells largely dependent on the physical interaction, and that these cytokines might be a contributing factor to the survival and proliferation of MDA-MB-231 when disseminated as heterotypic CTC-CAF clusters in the zebrafish.

## 4 Discussion

Current knowledge places CTCs, and in particular CTC-clusters, as key players in metastasis. Several studies have shown that CTC-clusters have a higher metastatic potential than individual CTCs (1, 8, 10), making these groups of cells important prognostic markers of metastatic disease (1, 2, 4, 6, 36–41). In addition, CTCs in clusters may not act alone, and they can be assisted by stromal cells from the tumour microenvironment (TME), such as immune cells, platelets, and cancer-associated fibroblasts (CAFs), whose presence in the clusters seem to enhance the metastatic potential of CTCs (30, 42–44). Indeed, analysis of the blood of cancer patients has shown the presence of circulating CAFs, associated in large proportion with CTCs in heterotypic CTC-clusters. This evidence manifests the importance of identifying the role that cells from the TME play within heterotypic CTC-clusters, and the mechanisms by which they may assist CTCs in metastasis seeding. In particular, preclinical and clinical evidence indicates that the interaction of CTCs with CAFs seems to exert an enhancement on the survival and metastatic potential of CTCs (24, 26, 30). CAFs are one of the most dominant cellular components in the TME, and although their role in tumourigenesis and promoting the dissemination of tumour cells is well established (15, 45, 46), little is yet known about the mechanism involved in CTC-CAF clustering and how they enhance CTC survival and metastases seeding.

CTC-clusters are found at a very low frequency in the blood of cancer patients making it difficult for their study. Thus, CTC-cluster models derived from functionally and molecularly diverse breast cancer cell lines, and their combination with stromal cells represent an opportunity to better study the biology of CTC-clusters and to deepen into the mechanisms that may shape their metastatic potential. In the present study, we have used BC models of homotypic and heterotypic clusters whose characterisation by functional and molecular assays has allowed us to understand the complexity of cellular interactions established in the clusters between the CTCs and CAFs. In particular, using a zebrafish model of metastasis, we show that the physical interactions established in the clustering of BC cells and CAFs seem to favour the selection of CTC populations with a higher capacity to survive and proliferate upon dissemination at the secondary sites. Importantly, this effect seems to be dependent on the maintained cell-cell contact between both cell types over time. While the main role attributed to CAFs as a population is to promote tumour progression and facilitate metastasis driving the invasion and dissemination of tumour cells, our results show that these cells can otherwise exert alternative regulatory effects on the CTCs without being involved in promoting tumour cell invasion. Such effects appear to be dependent on factors such as BC cell molecular phenotype, the crosstalk between tumour cells and CAFs (i.e. paracrine or physical), and the capacity of CAF to survive upon dissemination.

We show that BC cell lines MDA-MB-231 and MCF7 co-cultured or clustered with the vCAF fibroblasts display a different behavior in cellular processes associated with metastatic competence. Thus, while TNBC cells MDA-MB-231 showed a reduced capacity for proliferation, migration and colony formation when cocultured with vCAF, the effect on the luminal cells MCF7 was the opposite. Moreover, the magnitude of these effects was dependent on the fibroblast:BC cell ratio. These results indicate that highly invasive mesenchymal-like cells (due to an EMT) might compete with the fibroblast for energetic resources slowing their proliferation (47), while the epithelial tumour cells might possibly benefit from a fibroblast-directed migration (48) and survival signals, such as the secretion of extracellular matrix (ECM) components (49), alleviating cell death triggered under anchorage-independent conditions. Despite these results, the coculture of vCAFs with the BC cells was not able to induce significant molecular changes in genes (CDH1, CDH2, VIM) associated with epithelial-to-mesenchymal transition (EMT), a process mediating the acquisition of an invasive phenotype. In this regard, it is important to point out that the slight changes in gene expression observed could be translated into molecular changes at the protein level, and therefore functional, which we did not evaluate.

Despite the established role of CAFs at the primary site as promoters of tumour cell invasion and the higher tumour cell migration observed for heterotypic clusters in transwell assays, our zebrafish experiments showed that vCAF do not enhance the invasiveness of MDA-MB-231 cells when xenografted as heterotypic clusters. These clusters presented a similar ability to leave the injection site (primary tumour) as homotypic CTC-clusters and heterotypic clusters formed with normal fibroblasts (BJ). In line with this, the interactions in the clusters of vCAF and the non-metastatic MCF7 cells failed to induce a clear effect on CTC dissemination, but on the other hand, promoted the survival of tumour cells at the injection site. Therefore, our results suggest that some CAF populations may exert regulatory functions on BC cells other than promoting tumour cell invasion. Nevertheless, these data are at odds with previous studies in which various CAF populations have been shown to increase the metastatic capabilities of breast cancer cells when xenotransplanted into zebrafish and mouse models.(49–51)

The injection of heterotypic MDA-MB-231-vCAF clusters into the lateral tail vein of immunodeficient mice showed a delayed incidence of metastasis compared to homotypic CTC-clusters. This difference in metastasis formation between groups seems to be mainly established in the first 3 days upon injection, where the drop in bioluminescence signal was more pronounced for the heterotypic group of mice, suggesting that heterotypic clusters have a lower resistance to hemodynamic forces or a lower capacity for arrest and extravasate, or both, than the homotypic. Previous work with prostate cancer cell lines has reported that CAFs appear to be able to help tumour cells survive in circulation by conferring resistance to shear stress forces via intercellular contact and soluble derived factors (52). The *in vitro* FSS experiments, however, showed that the presence of the fibroblasts vCAF in the clusters did not have a significant effect on the survival and proliferation of the MDA-MB-231 cells, suggesting that the lower formation of lung metastases of heterotypic clusters is not due to an impaired resistance to shear stress in circulation. However, we cannot rule out the possibility that the massive tumour cell death experienced by heterotypic clusters is also due to severe necrosis caused by the death of CAFs, as these mice were injected with a higher total number of cells compared to mice injected with homotypic (5 x 10^5^ BC cells + 5 x 10^5^ CAF/mice vs. 5 x 10^5^ BC cells/mice). Importantly, histological examination of the lungs from mice injected with heterotypic clusters found no CAFs present in the tissues, suggesting that these cells did not survive. Together, these evidences seem to indicate that heterotypic CTC-clusters formed by MDA-MB-231 cells and vCAFs are less metastatic than homotypic clusters.

Later technological developments have revealed an important level of CAF heterogeneity existing in different tumour types, such as breast cancer, colorectal cancer and pancreatic cancer (53–57). Such heterogeneity could be an explanation for the findings of some reports showing that CAFs can also restrain tumour growth and progression (58, 59). Our *in vitro* assays and mice and zebrafish experiments comparing the behavior of homotypic and heterotypic CTC-clusters go in a similar direction, and present the vCAF fibroblasts as a break on the development of MDA-MB-231-mediated metastasis, with impaired cell proliferation, migration and colonisation of secondary sites, together with a non pro-invasive phenotype. Moreover, the similar gene expression profile of fibroblast activation markers (FAP, α-SMA and TNC) observed when comparing vCAF and BJ fibroblasts could be an explanation for the lack of CAF-mediate tumour dissemination enhancing phenotype, as activated fibroblasts expressing these markers are linked to tumour invasion and metastasis (29, 60, 61). This idea is further supported by the capacity of vCAFs to upregulate the expression of the markers, as well as to become pro-invasive population favouring BC CTC dissemination upon TGF-β stimulation. On the other hand, these results seem to suggest that the crosstalk established between CTCs and CAF plays an important role in the initial steps of metastasis.

Recent works suggest that CAFs may be involved in the formation of CTC-clusters and their metastatic success (15, 29). Our zebrafish model does not evaluate cluster formation, but instead, it allowed us to observe that the co-dissemination of CTCs and CAFs as heterotypic clusters is not due to a vCAF-driven invasion of MDA-MB-231 cells. Importantly, it also shows us that the successful homing of CTCs at the distant site is influenced by cell-to-cell interactions established with the CAFs in the cluster, which when preserved upon CTC-cluster arrest and extravasation, may promote CTC survival and proliferation. Remarkably, this effect was not observed when cell-to-cell interactions were established with the normal fibroblasts BJ (Figure 4C, F), nor when CTCs and CAFs were xenotransplanted as cell suspensions, without pre-existing cell-cell interactions (Figure 5B). Therefore, the physical communication between BC CTC and CAFs derived from cell clustering may represent a mechanism of fibroblast-mediated cancer metastasis, specific to CAFs and not to normal fibroblasts. In line with this, a report by Sharma et al., has shown the importance of CD44-mediated heterotypic clustering in a subset of CTCs and CAF in enhancing the metastatic capacity of BC CTCs (26). Moreover, a previous work using the zebrafish model has shown that disseminated CAFs can remain in close association with tumour cells at the metastatic foci formed, helping to promote the metastatic capacity of tumour cells (51). Similar results were observed in our analysis of CTC-CAF colocalization, which shows that this phenomenon – along with the accompanying proliferation of disseminated CTCs – is dependent on the survival of disseminated vCAFs. Therefore, the intercellular communication established in the clusters between CTCs and CAFs represents a survival mechanism for CTCs resulting in an enhanced metastatic capacity.

In support of this idea, we show that cytokines and chemokines released in cocultures of BC cells and vCAFs differ between cells in physical interaction and cells in paracrine interaction only. Thus, CCL5/RANTES, CXCL10/IP-10 and ICAM1/CD54 secretion are detected when MDA-MB-231 and vCAF cells are in physical communication, while CCL2/MCP1 is detected under paracrine conditions. High levels of CCL5 and CXCL10 in breast cancer correlate with advanced stages of the disease (62–65) and reduced survival of metastatic breast cancer patients (66), respectively. Besides, these cytokines can modulate the proliferation, motility, invasion and metastasis (33, 34). In particular, CXCL10 has been shown to regulate the proliferation and growth of MDA-MB-231 cell in metastatic niches in a metastasis model (66). Interestingly, ICAM1, a surface receptor activating pathways related to cell cycle and stemness, has been shown to mediate cell-cell interactions initiating the formation of CTC-clusters and favour CTC-cluster transendothelial migration and the formation of metastases (26, 35). Thus, our data suggest that these molecules may be contributing factors to the survival and proliferation of BC cells observed in zebrafish and, therefore, mediators of the metastatic potential of CTC-clusters.

Based on the above data, it is conceivable to speculate that vCAF might favour the selection of CTC subpopulations with a higher capacity to survive and proliferate upon dissemination. But we could also speculate that the survival of disseminated vCAF is representing a selection of CAF subpopulations harboring molecular and functional characteristics with the capacity to modify BC cell proliferation, due to the physical interaction between the two cell types. This hypothesis could be feasible in the light of the growing evidence showing CAF heterogeneity, in part due to the different origins of the CAFs responsible for their diversity within tumours (45, 46, 67, 68). However, further experiments should be performed to test such a hypothesis. Lastly, future work should be aimed at identifying the surface molecules mediating the clustering of CTC and CAF, as they may represent potential therapeutic targets.

## Supporting information

Hurtado et al Supplementary Material

## 6 Author Contributions

PH was involved in methodology, data analysis and the original draft preparation. IM-P, SY-R, MB-O, and CF-S were involved in methodology, and data analysis. CA was involved in methodology. LS was involved in the methodology and interpretation of the data. RL-L was involved in the design and conceptualization of the study, interpretation of the data and drafting and revision of the manuscript. RP was involved in design and conceptualization of the study, supervision, data analysis, interpretation of the data and drafting and revision of the manuscript.

## 8 Acknowledgments

We kindly thank Prof. Erik Sahai for providing the vCAF cells. We thank the personnel from the Experimental Biomedicine Centre (CEBEGA) of the University of Santiago de Compostela for their technical support. We thank Jose Antonio Trillo Franco for his technical assistance with the *in vitro* experiemnts.

