## Supplementary material for "Modelling metastasis in zebrafish unveils regulatory interactions of cancer-associated fibroblasts with circulating tumour cells": Hurtado et al Supplementary Material

### 1. Supplementary Figures

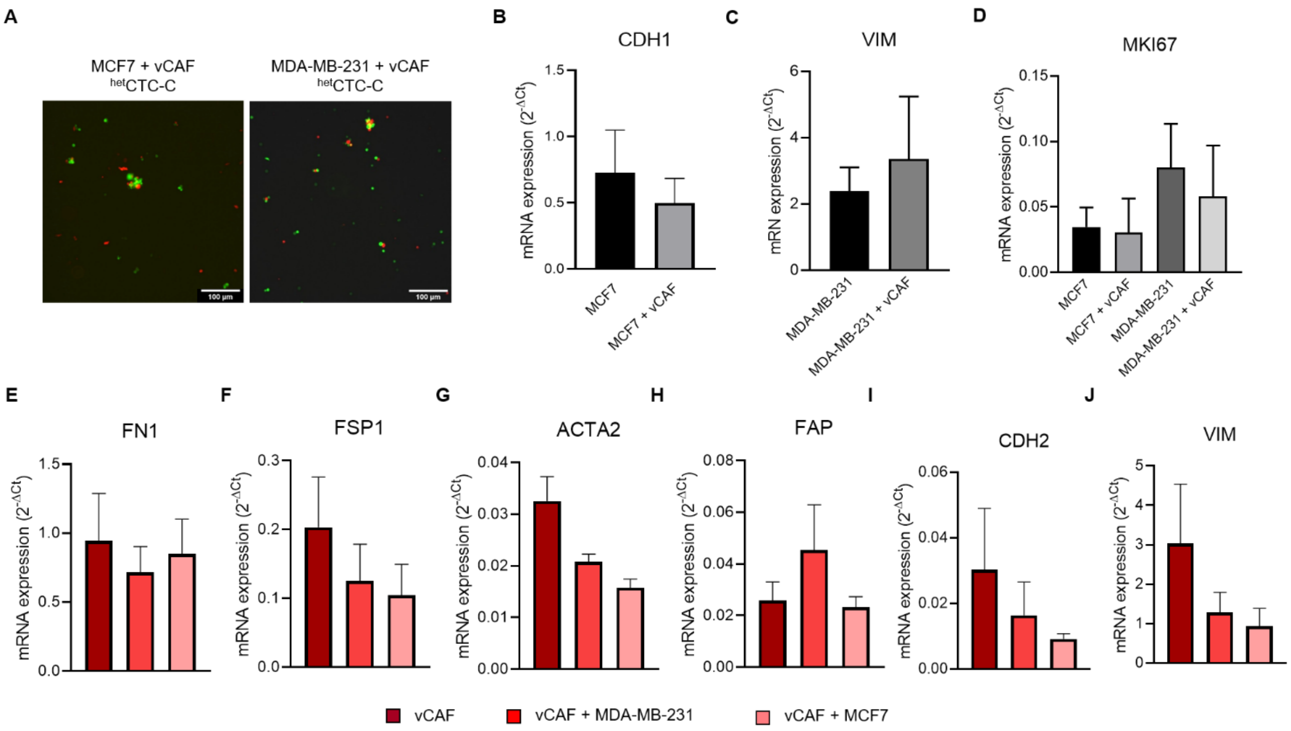


**Supplementary Figure 1.** Gene expression analysis of BC cell lines and vCAF under paracrine interactions in co-cultures. **(A)** Representative images of heterotypic clusters formed by BC cells MCF7 or MDA-MB-231 in combination with the fibroblasts vCAF; Relative mRNA expression observed in BC cell monocultures and co-cultures for the genes CDH1 **(B)**, VIM **(C)**, and MKI67 **(D)**; **(E-J)** Relative mRNA expression detected in vCAF in monoculture and co-culture with BC cells for mesenchymal and CAF markers. Data is expressed as $2^{-\Delta CT}$, relative to the average expression levels of β-2-microglobulin (β2M), and Glyceraldehyde-3-Phosphate Dehydrogenase (GAPDH), which were used as the housekeeping genes.

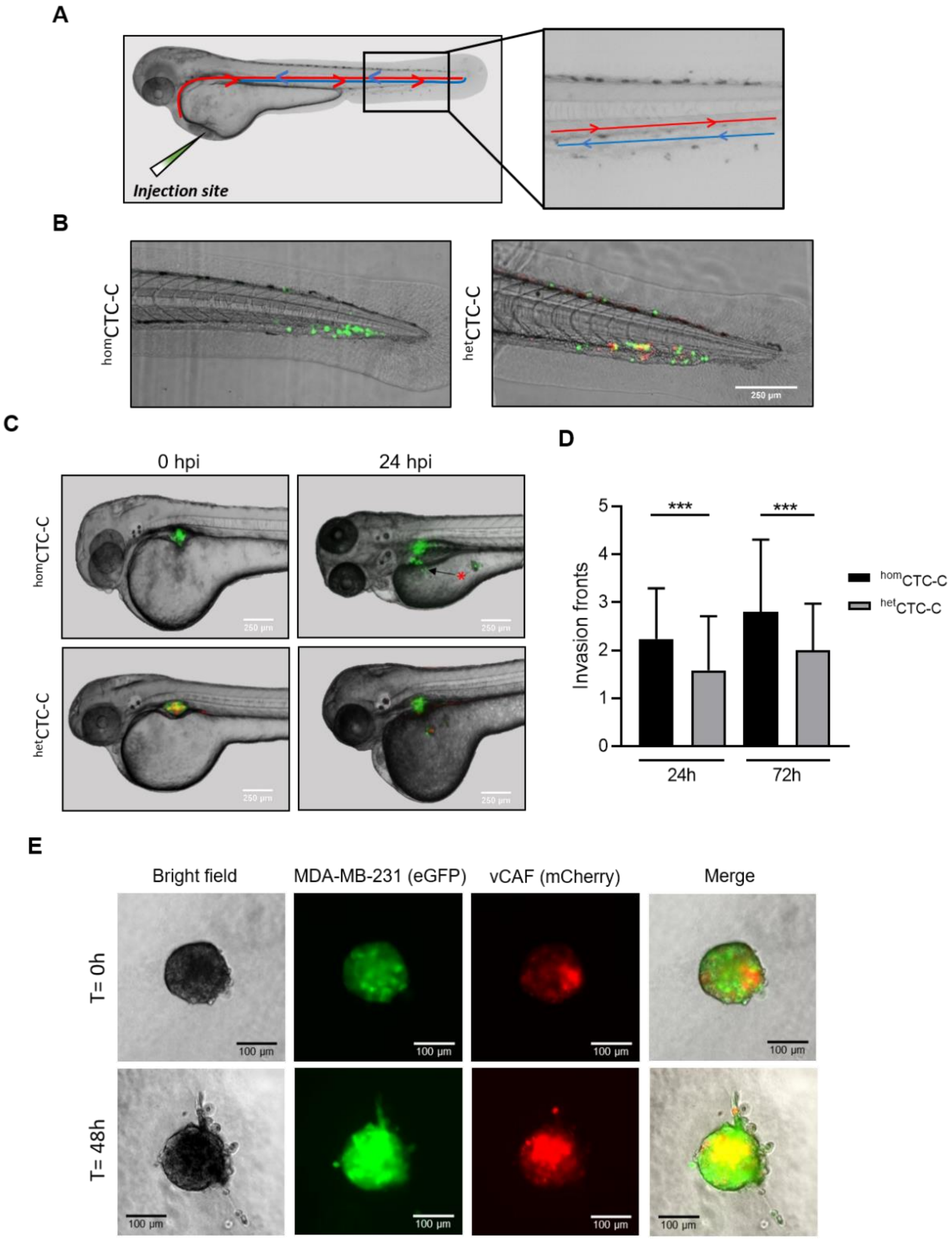


**Supplementary Figure 2.** Characterisation of MDA-MB-231 homCTC-C and hetCTC-C in the zebrafish embryo. **(A)** Schematic representation of the zebrafish embryo circulation and the location of cell dissemination at the caudal region; **(B)** Representative images of disseminated cells in the tails of homCTC-C and hetCTC-C xenografted fish at 72 hours post-injection (hpi) (scale bar 250µm); **(C)** Representative images of zebrafish showing homCTC-C and hetCTC-C from MDA-MB-231 injected at the perivitelline space right after injection (0 hpi) and 24 hpi. The asterisk marks the presence of an invasion front (scale bar 250µm); **(D)** Quantification of invasion fronts generated by MDA-MB-231 homCTC-C and hetCTC-C at 24-72 hpi; **(E)** Representative images of the invasive pattern of spheroids formed by eGFP-labelled MDA-MB-231 (green) and mCherry-labelled vCAF. (***$p < 0.001$).

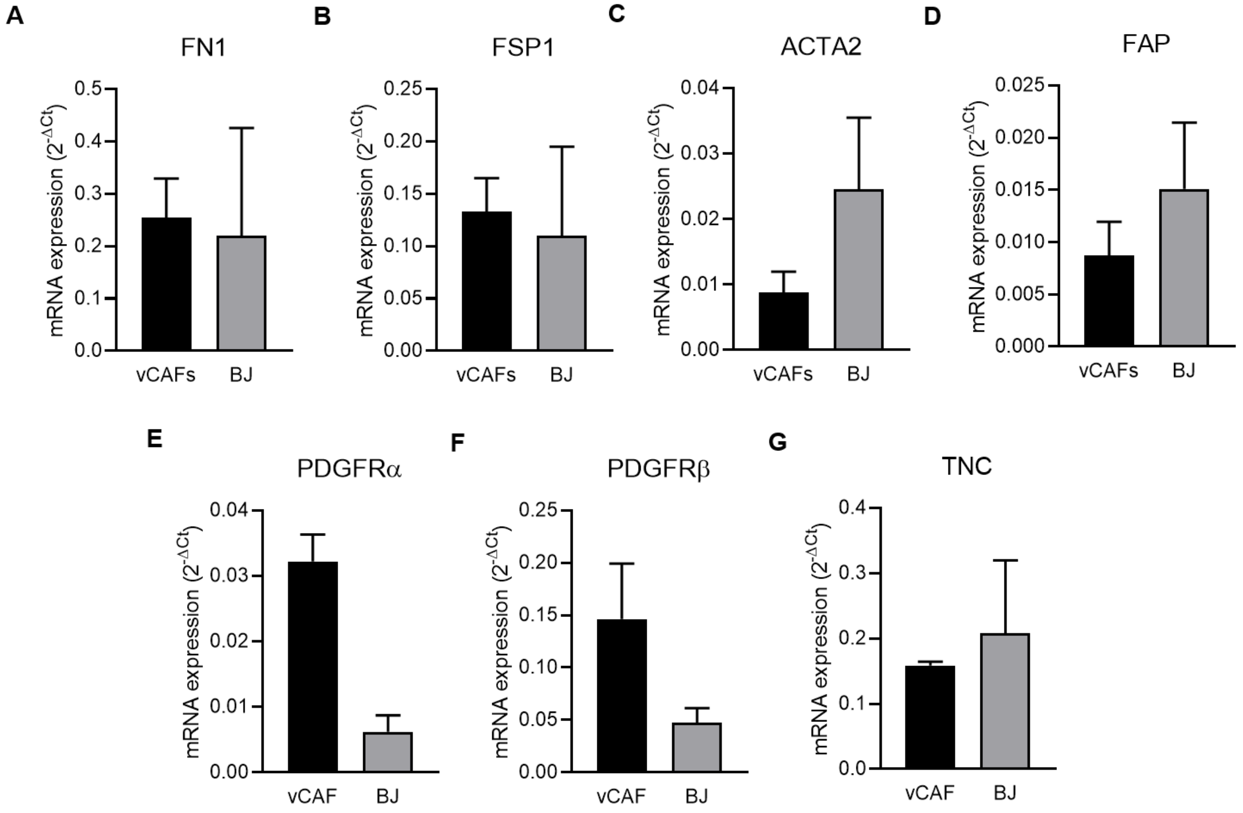


**Supplementary Figure 3.** Comparative gene expression analysis of fibroblast activation markers between vCAF and BJ. Relative mRNA expression levels detected in the fibroblasts for the genes **(A)** FN1, **(B)** FSP1, **(C)** ACTA2, **(D)** FAP, **(E)** PDGFRα, **(F)** PDGFRβ, and **(G)** TNC. Data are expressed as $2^{-\Delta CT}$, relative to the average expression levels of β-2-microglobulin (β2M), and Glyceraldehyde-3-Phosphate Dehydrogenase (GAPDH), which were used as the housekeeping genes (n= 3).

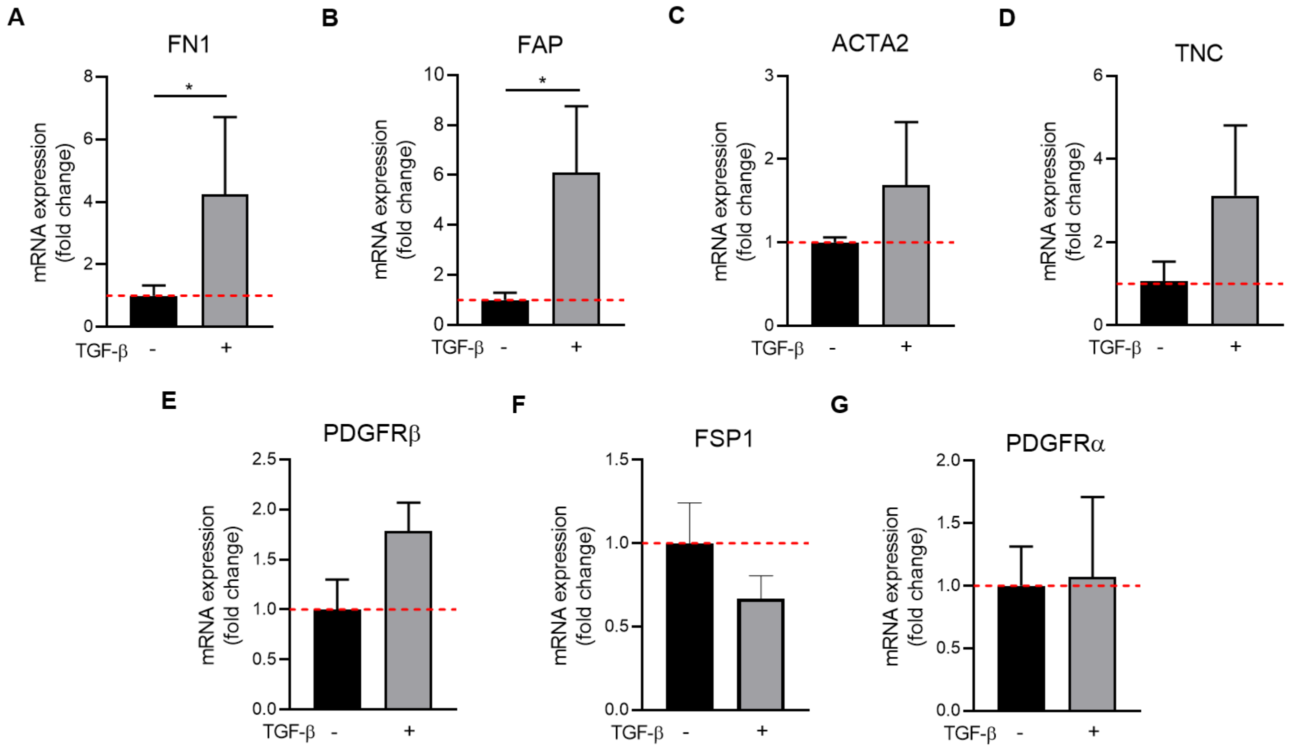


**Supplementary Figure 4.** Effect of TGF-β treatment in the gene expression levels of fibroblast activation markers in fibroblasts vCAF. Relative mRNA expression levels detected in the fibroblasts vCAF stimulated or not with 5ng/ml TGF-β for 48 hours for the genes **(A)** FN1, **(B)** FAP, **(C)** ACTA2, **(D)** TNC, **(E)** PDGFRβ, **(F)** FSP1, and **(G)** PDGFRα. Data are expressed as fold change of the $2^{-\Delta CT}$ between treated and untreated cells. $2^{-\Delta CT}$ is calculated relative to the average expression levels of β2M and GAPDH (n= 4). (*$p < 0.05$).

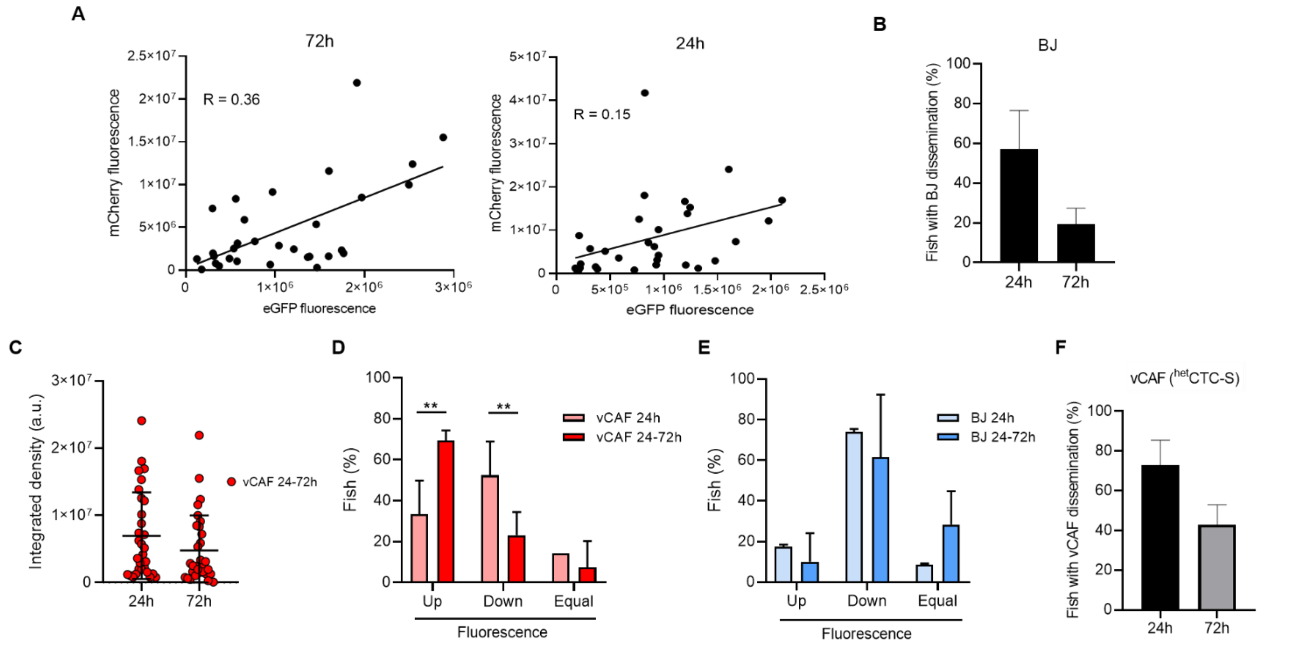


**Supplementary Figure 5.** Analysis of the dissemination and survival of MDA-MB-231 and vCAF or BJ $^{het}$CTC-C in the zebrafish embryo. **(A)** Linear regression analysis of the fluorescence emitted by MDA-MB-231 (eGFP) and vCAF (mCherry) cells disseminated in zebrafish tails at 24 and 72 hpi; **(B)** Percentage of fish xenografted with MDA-MB-231 and BJ $^{het}$CTC-C showing disseminated BJ in the tail at 24 and 72 hpi; **(C)** Dissemination volume of vCAF at 24 and 72 hpi in fish whose fibroblasts survive at 24 and 72 hours (vCAF 24-72) based on the integrated density of the fluorescence emitted by mCherry; **(D)** Percentage of fish xenografted with $^{hom}$CTC-C and vCAF $^{het}$CTC-C whose fluorescence in the tail has increased, decreased, or maintained over time (n =2 independent experiments); **(E)** Percentage of fish xenografted with $^{hom}$CTC-C and BJ $^{het}$CTC-C whose fluorescence in the tail has increased, decreased, or maintained over time (n =3 independent experiments); **(F)** Percentage of fish xenografted with MDA-MB-231 and vCAF $^{het}$CTC-S showing disseminated fibroblasts in the tail at 24 and 72 hpi. (***p* < 0.01).

2. **Supplementary Tables**

**Supplementary Table 1.** Panel of genes analyzed by RT-qPCR using Taqman probes

| Gene | Taqman Assay |
|---|---|
| ACTA2 | Hs00426835_g1 |
| B2M | Hs00187842_m1 |
| CD44 | Hs01075861_m1 |
| CDH1 | Hs00170423_m1 |
| CDH2 | Hs00983056_m1 |
| FAP | Hs00990806_m1 |
| FN1 | Hs01549976_m1 |
| FSP1 | Hs00243202_m1 |
| GAPDH | Hs99999905_m1 |
| MKI67 | Hs01032443_m1 |
| PDGFRA | Hs00998018_m1 |
| PDGFRB | Hs01019589_m1 |
| TNC | Hs01115665_m1 |
| VIM | Hs00958116_m1 |

**Supplementary Table 2.** Panel of cytokines/chemokines analyzed by the cytokine array

| Target | Entrez Gene ID |
|---|---|
| CCL1/I-309 | 6346 |
| CCL2/MCP-1 | 6347 |
| MIP-1α/MIP-1β | 6348/6351 |
| CCL5/RANTES | 6352 |
| CD40 | 959 |
| Complement | 727 |
| CXCL1/GROα | 2919 |
| CXCL10/IP-10 | 3627 |
| CXCL11/I-TAC | 6373 |
| CXCL12/SDF-1 | 6387 |
| G-CSF | 1440 |
| GM-CSF | 1437 |
| ICAM-1/CD54 | 3383 |
| IFN-γ | 3458 |
| IL-1α/IL-1F1 | 3552 |
| IL-1β/IL-1F2 | 3553 |
| IL-1ra/IL-1F3 | 3557 |
| IL-2 | 3558 |
| IL-4 | 3565 |
| IL-5 | 3567 |
| IL-6 | 3569 |

| | |
|---|---|
| IL-8 | 3576 |
| IL-10 | 3586 |
| IL-12 p70 | 3592/3593 |
| IL-13 | 3592/3593 |
| IL-16 | 3603 |
| IL-17A | 3605 |
| IL-17E | 3605 |
| IL-18/IL-1F4 | 3606 |
| IL-21 | 59067 |
| IL-27 | 246778 |
| IL-32α | 9235 |
| MIF | 4282 |
| Serpin E1/PAI-1 | 5054 |
| TNF-α | 7124 |
| TREM-1 | 54210 |
